# Cohort Specific Effects of Cereal-bar Supplementation in Overweight Patients With or Without Type 2 Diabetes Mellitus

**DOI:** 10.1101/066704

**Authors:** Chieh Jason Chou, Chris Lauber, Anirikh Chakrabarti, Jay Siddharth, Anne Chalut-Carpentier, Zoltan Pataky, Alain Golay, Scott Parkinson

**Affiliations:** Nestlé Institute of Health Sciences S.A., Lausanne, Switzerland; Service of Therapeutic Education for Chronic Diseases, WHO Collaborating Centre, University Hospitals of Geneva and University of Geneva, Geneva, Switzerland

## Abstract

The importance of gut microbes to metabolic health is becoming more evident and nutrition-based therapies to alter the composition of bacterial communities to manage metabolic disease are an attractive avenue to ameliorate some effects of Western diets. While the composition of gut microbial communities can vary significantly across disease states, it is not well known if these communities have common responses to nutritional interventions. To better understand fiber-bacterial community interactions, we collected biological parameters and fecal samples of overweight non-diabetic (OND) and diabetic (OD) individuals before and after daily supplementation of 2.8 g β-glucan on their habitual diet for 30 days. Fecal bacterial communities in an age-matched cohort were measured by sequencing partial 16S rRNA genes and imputed metagenomic content. Unexpectedly, we observed disconnected responses of biological measurements and the bacterial community. Based on average effect size, biological measurements were greater in the OND group while effects on the bacterial community were greatest on the OD cohort, and we suspect these observations are due to the significantly lower alpha diversity in the OD cohort. Our data indicate that responses to fiber supplementation are cohort specific and this should be considered when manipulating the microbiome via fiber supplementation.

## Introduction

As a result of a globalization of the economy, obesity and diabetes are becoming major issues not just for the developed but the developing world. The global adoption of a Western lifestyle includes dietary modification to diets low in fiber and high energy density. Deaths accounted for by non-communicable diseases have now overtaken deaths due to communicable diseases virtually world wide [1]. This statistic highlights the dilemma for providing high quality fiber rich foods and avoiding over nutrition for both the developed and developing world.

The gut bacterial community serves as one conduit by which the consequences of nutrition and dietary choices integrate into human health, and therefore must be considered in the context of any nutritional intervention. Recent work shows abundances of dominant taxa in the distal gut vary widely across individuals and habitual food choices [2-6]. As Firmicutes and Bacteroidetes are by far the most prevalent taxa in the human gastro-intestinal tract [7], their presence and response to diet can have a significant impact on human health. For instance, members of the *Fecalbacterium* (a Firmicute) produce short-chain fatty acids such as butyrate [8] that serve as an energy source for colonic epithelia. *Bacteroides and Ruminococcaceae* (both Bacteroidetes) degrade polysaccharides found in foods containing fiber [9, 10]. Conversely, abundant taxa such as the Proteobacteria are typically associated with dysbiosis observed in persons with inflammatory bowel disorders [11, 12] or in the guts of those consuming a high fat or typical Western style diet [2]. Despite the inter- and intra-individual variability in the abundance of bacterial taxa, these and other studies demonstrate the importance of the bacterial communities to integrate diet and host metabolism. Dietary supplements aiming to leverage the benefits of microbial function therefore may become practical solutions to Western lifestyle associated diseases if their efficacy and the molecular mechanisms underlying the bacteria-diet interactions could be demonstrated.

Fiber supplementation in the diet is one approach proposed to minimize the effects of Western life styles by potentially altering the composition and metabolism of the gut bacterial community [13, 14]. However it is not widely known if fiber has the same effect on microbiomes in subjects of related but distinct disease states. For instance, there are numerous studies that document fiber induced alterations in the microbiomes of monotonic cohorts [15-19]. Hooda et al. [20] demonstrated the significant effect of corn derived fibers on abundances of genera of belonging to Firmicutes, Bacteroidetes and Verrucomicrobia in generally healthy subjects, while Benus et al. [21] further demonstrated that pea fiber and fructo-oligosaccharides supplementation was associated with increased abundances of butyrate producing bacteria in healthy subjects. Though these studies have given us great insight into the effect of fiber on the microbiome, each used unique diet amendments in cohorts that consisted almost exclusively of healthy, overweight, obese, or subjects with diagnosed metabolic disease. Given that bacterial community composition and diversity vary significantly across disease states, it is reasonable to assume that effects in one cohort may not be applicable to others. This lack of knowledge may be important when identifying patients whose microbiomes may be more or less responsive to fiber supplementation and thus resistant to improvement of host metabolic markers potentially mediated by gut bacteria. Thus quantifying the responsiveness of the microbiome in similar but distinctive cohorts may help to define the limits of fiber supplementation via changes in the microbiome.

Overweight diabetic (OD) and overweight non-diabetics (OND) represent two metabolic states that can be distinguished by the degree of insulin resistance. However, it is not known whether the two populations exhibit cohort specific effects of fiber supplementation on the bacterial communities and host metabolism. To answer this question, we conducted a fiber supplementation study in OD and OND subjects with a cereal bar containing 1.4 g β-glucan. We investigated how the bacterial communities responded to consumption of two cereal bars per day and determined whether there were unique features of the response in OND vs OD subjects. Specifically we predicted that fiber supplementation would increase abundances of Bacteroidetes and Verrucomicrobia while decreasing Firmicutes and Proteobacteria in both cohorts with concomitant improvement in host metabolic markers.

## Results

### Biological Parameters

We recruited 46 overweight and obese individuals with and without diabetes to assess the effect of a β-glucan containing cereal bar supplementation on biological parameters and the distal gut bacterial community. The subject characteristics are summarized in S1 Table. Subjects went through a diet normalization phase followed by consumption of 2 cereal bars per day for 30 days to deliver 2.8g β-glucan fiber to the habitual diet. Twenty-six individuals were excluded for compliance issues, lack of sequencing success, or fell outside the age-range, leaving 10 individuals in each group for analysis. Non-parametric testing indicated no significant difference in any biological measurement between the cohorts at the pre-supplementation time point. However, LDL and triglyceride concentrations were statistically different between the OND and OD post supplementation (Table 1, S2 Table), though these differences were not significant improvements from their pre-supplementation values. The remaining biological parameters were unaffected by consumption of the cereal bar. Lastly, Kruskal-Wallis tests showed there were no differences within each cohort between pre- and post-supplementation time points (Table1).

**S1 Table.** Mean and standard deviation of biological parameters of 46 subjects at recruitment.

|  | OND | OD |
| --- | --- | --- |
| Subjects | 25 | 21 |
| Height(cm) | 166(8.7) | 169(9.5) |
| Age | 43(7.2) | 50(5.9) |
| Weight(kg) | 91(14) | 95(16) |
| BMI (kg m<sup>2</sup>) | 33(4.8) | 33(4.3) |
| WaistCircumference(cm) | 102(11) | 112(11) |
| HipCircumference(cm) | 116(9.1) | 113(9.5) |
| Diastolic BP (mmHg) | 89(13) | 87(11) |
| Systolic BP (mmHg) | 124(15) | 130(16) |
| Cardiac Frequency (beats min<sup>-1</sup>) | 73(11) | 81(19) |
| Total Cholesterol (mmol L<sup>-1</sup>) | 5.0(0.8) | 4.7(1.3) |
| Triglyceride (mmol L<sup>-1</sup>) | 1.1(0.4) | 2.9(4.2) |
| HDL cholesterol (mmol L<sup>-1</sup>) | 1.1(0.2) | 0.9(0.2) |
| LDL (mmol L<sup>-1</sup>) | 3.4(0.7) | 2.8(0.9) |
| Glucose (mmol L<sup>-1</sup>) | 4.9(0.6) | 6.9(1.9) |
| Insulin (uIU ml<sup>-1</sup>) | 18(11) | 23(19) |
| HOMA_IR | 4.0(2.8) | 7.4(6.2) |
| LBP (ug ml<sup>-1</sup>) | 25(8) | 29(7.6) |
| FFA (mmol L<sup>-1</sup>) | 0.45(0.21) | 0.54(0.21) |

**Table 1.** Biological parameters of age-matched cohorts at enrollment, pre-supplementation, and post-supplementation. The mean and standard deviation (in parentheses) are shown. Except for *Age and *Height, the rest of parameters were included in biological effect size calculation. Superscript “a” indicates significant difference between cohorts post-supplementation.

|  | OND Mean |  |  | OD Mean |  |  |
| --- | --- | --- | --- | --- | --- | --- |
|  | Enrollment | Pre-supplementation | Post-supplementation | Enrollment | Pre-supplementation | Post-supplementation |
| *Age | 48 (3.5) | 48 (3.5) | 48 (3.5) | 47 (4.3) | 47 (4.3) | 47 (4.3) |
| *Height (cm) | 167 (10.3) | 167 (10.3) | 167 (10.3) | 171 (10) | 171 (10) | 171 (10) |
| Weight (kg) | 95 (14.7) | 95 (14.1) | 94 (14.4) | 98 (17) | 96 (17) | 96 (16) |
| BMI (kg m<sup>-2</sup>) | 34 (5.2) | 34 (5.2) | 34 (5.3) | 34 (4.66) | 33 (4.5) | 33 (4.4) |
| Waist Circumference (cm) | 104 (11.4) | 105 (11.3) | 106 (11.3) | 113 (10) | 113 (10) | 113 (12) |
| Hip Circumference (cm) | 117 (8.1) | 117 (8.3) | 116 (8.2) | 116 (12) | 114 (11) | 115 (12) |
| Diastolic BP (mmHg) | 95 (16) | 87 (5.1) | 89 (13) | 87 (9.86) | 85 (9.63) | 88 (14) |
| Systolic BP (mmHg) | 131 (15) | 124 (13) | 131 (21) | 130 (17) | 129 (22) | 131 (17) |
| Cardiac Frequency (beats min<sup>-1</sup>) | 67 (11) | 68 (12) | 72 (16) | 79 (24) | 78 (13) | 77 (14) |
| Total Cholesterol (mmol L<sup>-1</sup>) | 5 (0.74) | 4.9 (0.87) | 5.2 (0.69) | 4.9 (1.3) | 4.6 (0.82) | 4.7 (0.49) |
| Triglyceride (mmol L<sup>-1</sup>) | 1.1 (0.42) | 1.2 (0.51) | 1.2 (0.43) | 4.1 (5.8) | 3.6 (4.3) | 2.9 (2.1) |
| HDL cholesterol (mmol L<sup>-1</sup>) | 1 (0.23) | 1 (0.3) | 1.1 (0.25) | 0.9 (0.24) | 0.8 (0.24) | 0.8 (0.16) |
| LDL (mmol L<sup>-1</sup>) | 3.5 (0.61) | 3.4 (0.68) | 3.6 (0.65)<sup>a</sup> | 2.8 (0.74) | 2.7 (0.7) | 2.7 (0.67)<sup>a</sup> |
| Glucose (mmol L<sup>-1</sup>) | 5 (0.68) | 5 (0.63) | 5.1 (0.46)<sup>a</sup> | 6.4 (2) | 6.5 (2.2) | 6.3 (1.5)<sup>a</sup> |
| Insulin (uIU ml<sup>-1</sup>) | 17 (7.1) | 19 (8.1) | 19 (7.6) | 20 (13) | 17 (10) | 18 (12) |
| FFA (mmol L<sup>-1</sup>) | 0.5 (0.13) | 0.5 (0.2) | 0.6 (0.23) | 0.6 (0.25) | 0.5 (0.19) | 0.5 (0.19) |
| HOMA_IR | 3.9 (1.8) | 4.4 (2.1) | 4.3 (1.6) | 6.2 (5.8) | 5.1 (3.7) | 5.5 (5) |
| LBP (ug ml<sup>-1</sup>) | 28 (6) | 27 (3.8) | 29 (4.8) | 31 (8.7) | 27 (7.4) | 29 (7.7) |

**S2 Table.** Means, standard deviations, and effect sizes for biological parameters for the age-matched cohort. *Age and *height are not included in the mean effect size calculation. Superscript “a” indicates significant difference between cohorts post-supplementation.

|  | OND Mean |  |  | OD Mean |  |  | OND Effect Size |  | OD Effect Size |  |
| --- | --- | --- | --- | --- | --- | --- | --- | --- | --- | --- |
|  | Enrollment | Pre-supplementation | Post-supplementation | Enrollment | Pre-supplementation | Post-supplementation | Enrolment vs Pre-supplementation | Pre-supplementation vs Post-supplementation | Enrolment vs Pre-supplementation | Pre-supplementation vs Post-supplementation |
| Biological Parameters |  |  |  |  |  |  |  |  |  |  |
| *Age | 48 (3.5) | 48 (3.5) | 48 (3.5) | 47 (4.3) | 47 (4.3) | 47 (4.3) | 0.0000 | 0.0000 | 0.0000 | 0.0000 |
| *Height (cm) | 167 (10.3) | 167 (10.3) | 167 (10.3) | 171 (10) | 171 (10) | 171 (10) | 0.0000 | 0.0000 | 0.0000 | 0.0000 |
| Weight (kg) | 95 (14.7) | 95 (14.1) | 94 (14.4) | 98 (17) | 96 (17) | 96 (16) | 0.0007 | 0.0653 | 0.0873 | 0.0356 |
| BMI (kg m<sup>-2</sup>) | 34 (5.2) | 34 (5.2) | 34 (5.3) | 34 (4.66) | 33 (4.5) | 33 (4.4) | 0.0046 | 0.0610 | 0.1135 | 0.0427 |
| Waist Circumference (cm) | 104 (11.4) | 105 (11.3) | 106 (11.3) | 113 (10) | 113 (10) | 113 (12) | 0.0746 | 0.0664 | 0.0529 | 0.0091 |
| Hip Circumference (cm) | 117 (8.1) | 117 (8.3) | 116 (8.2) | 116 (12) | 114 (11) | 115 (12) | 0.0000 | 0.1698 | 0.1457 | 0.1008 |
| Diastolic BP (mmHg) | 95 (16) | 87 (5.1) | 89 (13) | 87 (9.86) | 85 (9.63) | 88 (14) | 0.6578 | 0.1860 | 0.2156 | 0.3138 |
| Systolic BP (mmHg) | 131 (15) | 124 (13) | 131 (21) | 130 (17) | 129 (22) | 131 (17) | 0.4325 | 0.3575 | 0.0502 | 0.1059 |
| Cardiac Frequency (beats min<sup>-1</sup>) | 67 (11) | 68 (12) | 72 (16) | 79 (24) | 78 (13) | 77 (14) | 0.1277 | 0.2319 | 0.0731 | 0.0223 |
| Total Cholesterol (mmol L<sup>-1</sup>) | 5 (0.74) | 4.9 (0.87) | 5.2 (0.69) | 4.9 (1.3) | 4.6 (0.82) | 4.7 (0.49) | 0.1384 | 0.3816 | 0.3066 | 0.1585 |
| Triglyceride (mmol L<sup>-1</sup>) | 1.1 (0.42) | 1.2 (0.51) | 1.2 (0.43) | 4.1 (5.8) | 3.6 (4.3) | 2.9 (2.1) | 0.0864 | 0.0352 | 0.1046 | 0.1999 |
| HDL cholesterol (mmol L<sup>-1</sup>) | 1 (0.23) | 1 (0.3) | 1.1 (0.25) | 0.9 (0.24) | 0.8 (0.24) | 0.8 (0.16) | 0.0224 | 0.1082 | 0.0955 | 0.0025 |
| LDL (mmol L<sup>-1</sup>) | 3.5 (0.61) | 3.4 (0.68) | 3.6 (0.65) | 2.8 (0.74) | 2.7 (0.7) | 2.7 (0.67) | 0.2127 | 0.4162 | 0.0680 | 0.0038 |
| Glucose (mmol L<sup>-1</sup>) | 5 (0.68) | 5 (0.63) | 5.1 (0.46) | 6.4 (2) | 6.5 (2.2) | 6.3 (1.5) | 0.0381 | 0.2262 | 0.0402 | 0.0813 |
| Insulin (uIU ml<sup>-1</sup>) | 17 (7.1) | 19 (8.1) | 19 (7.6) | 20 (13) | 17 (10) | 18 (12) | 0.3149 | 0.0406 | 0.1913 | 0.0487 |
| FFA (mmol L<sup>-1</sup>) | 0.5 (0.13) | 0.5 (0.2) | 0.6 (0.23) | 0.6 (0.25) | 0.5 (0.19) | 0.5 (0.19) | 0.1759 | 0.3637 | 0.4917 | 0.0354 |
| HOMA_IR | 3.9 (1.8) | 4.4 (2.1) | 4.3 (1.6) | 6.2 (5.8) | 5.1 (3.7) | 5.5 (5) | 0.2842 | 0.0476 | 0.2236 | 0.0908 |
| LBP (ug ml<sup>-1</sup>) | 28 (6) | 27 (3.8) | 29 (4.8) | 31 (8.7) | 27 (7.4) | 29 (7.7) | 0.1829 | 0.3826 | 0.4042 | 0.1908 |
| Biological Mean Effect Size |  |  |  |  |  |  | 0.1721 | 0.1962 | 0.1665 | 0.0901 |

### Taxon abundance and KEGG functions pre- and post-supplementation

Bacterial communities in each cohort were dominated by Firmicutes and Bacteroidetes, followed by the Actinobacteria, Proteobacteria and Verrucomicrobia (Fig 1A-1D, S3 Table). Abundances at the pre-supplementation time point were, however, not statistically different between OND and OD groups (Fig 1A-1D). Likewise the abundances of KEGG functions from predicted metagenomes were not different between the cohorts pre-supplementation (S4 Table). However, we did observe a significant difference in alpha diversity between OND and OD groups at enrollment and before the intervention (Fig 1E, S3 Table). The OND cohorts had significantly greater community richness compared to the OD group (Fig 1E, S3 Table).

**Fig 1.**
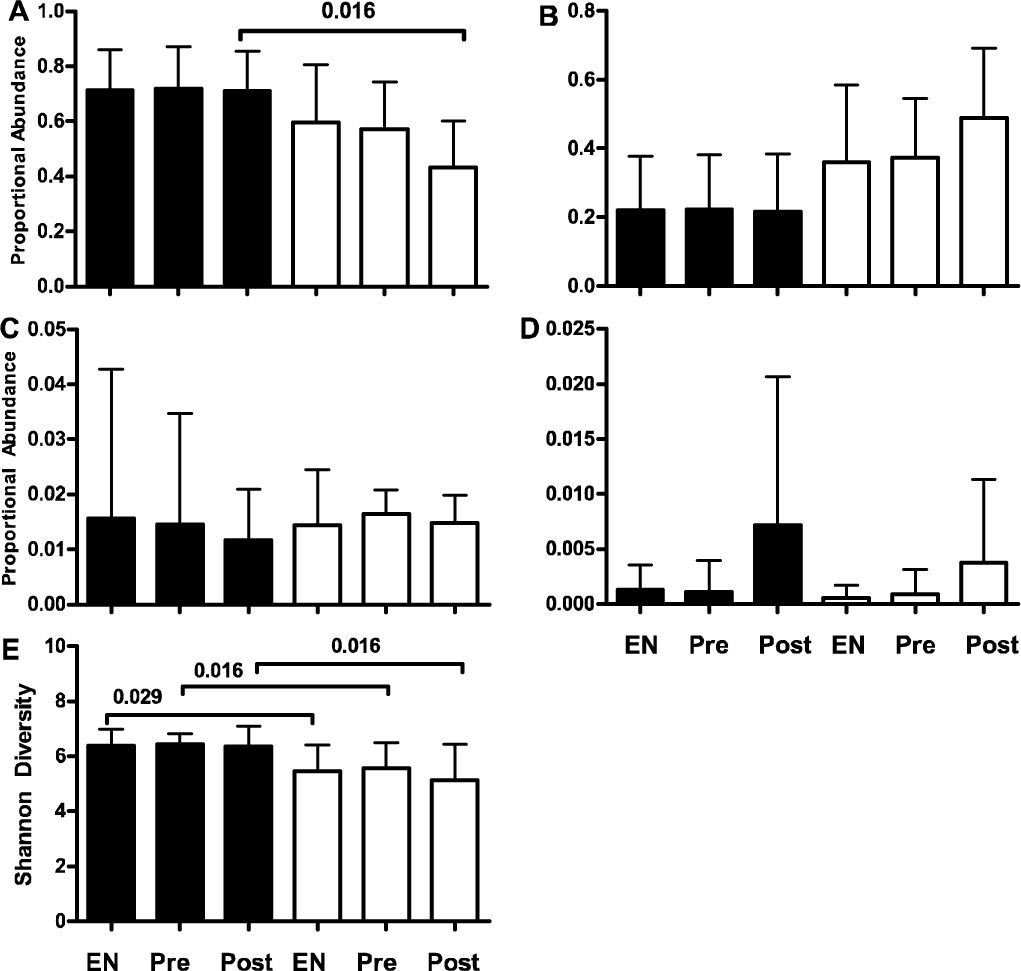
Proportional abundances of dominant taxa and Shannon diversity at enrollment (EN), pre-supplementation (Pre) and post-supplementation (Post). Bar graphs for proportional abundances of dominant taxa are indicated as (A) Firmicutes, (B) Bacteroidetes, (C) Proteobacteria, and (D) Verrucomicrobia. Filled bars represent the OND cohort, open bars represent the OD group. Error bars depict the standard deviation for the mean of 10 subjects. p-values for significant differences are shown.

**S3 Table.** Means, standard deviations, and effect sizes for taxa and Shannon Diversity for the age-matched cohort.

|  | OND Mean |  |  | OD Mean |  |  | OND Effect Size |  | OD Effect Size |  |
| --- | --- | --- | --- | --- | --- | --- | --- | --- | --- | --- |
|  | Enrollment | Pre-supplementation | Post-supplementation | Enrollment | Pre-supplementation | Post-supplementation | Enrolment vs Pre-supplementation | Pre-supplementation vs Post-supplementation | Enrolment vs Pre-supplementation | Pre-supplementation vs Post-supplementation |
| Taxa |  |  |  |  |  |  |  |  |  |  |
| Euryarchaeota | 0.003 (0.004) | 0.001 (0.002) | 0.003 (0.006) | 0 (0) | 0 (0) | 0 (0) | 0.3417 | 0.2984 | 0.1805 | 0.2959 |
| Actinobacteria | 0.043 (0.049) | 0.039 (0.053) | 0.048 (0.074) | 0.028 (0.04) | 0.036 (0.064) | 0.059 (0.096) | 0.0670 | 0.1410 | 0.1557 | 0.2819 |
| Bacteroidetes | 0.221 (0.157) | 0.222 (0.16) | 0.216 (0.169) | 0.361 (0.225) | 0.373 (0.172) | 0.49 (0.202) | 0.0100 | 0.0378 | 0.0606 | 0.6194 |
| Firmicutes | 0.714 (0.145) | 0.718 (0.154) | 0.71 (0.145) | 0.596 (0.209) | 0.572 (0.171) | 0.432 (0.17) | 0.0275 | 0.0573 | 0.1265 | 0.8221 |
| Fusobacteria | 0 (0) | 0 (0) | 0 (0) | 0 (0.001) | 0.002 (0.006) | 0 (0) | 0.6708 | 0.5855 | 0.3444 | 0.4316 |
| Lentisphaerae | 0 (0.001) | 0 (0) | 0 (0.001) | 0 (0) | 0 (0) | 0 (0) | 0.3087 | 0.2211 | 0.4884 | 0.6174 |
| Proteobacteria | 0.016 (0.027) | 0.015 (0.02) | 0.012 (0.009) | 0.014 (0.01) | 0.016 (0.004) | 0.015 (0.005) | 0.0405 | 0.1907 | 0.2602 | 0.3567 |
| Tenericutes | 0.002 (0.003) | 0.003 (0.003) | 0.004 (0.007) | 0 (0) | 0 (0) | 0.001 (0.003) | 0.0697 | 0.2273 | 0.4346 | 0.4626 |
| Verrucomicrobia | 0.00126 (0.002) | 0.001118 (0.003) | 0.00716 (0.014) | 0.001 (0.001) | 0.001 (0.002) | 0.004 (0.008) | 0.0549 | 0.6187 | 0.1758 | 0.5085 |
| Phyla Mean Effect Size |  |  |  |  |  |  | 0.1768 | 0.2642 | 0.2474 | 0.4885 |
| Richness |  |  |  |  |  |  |  |  |  |  |
| Shannon Index | 6.4 (0.60) | 6.4 (0.35) | 6.4 (0.74) | 5.5 (0.9) | 5.6 (0.9) | 5.1 (1.3) |  |  |  |  |

**S4 Table.** Means, standard deviations, and effect sizes for the KEGG functions predicted by PiCRUSt for the age-matched cohort.

|  | OND Mean |  |  | OD Mean |  |  | OND Effect Size |  | OD Effect Size |  |
| --- | --- | --- | --- | --- | --- | --- | --- | --- | --- | --- |
|  | Enrollment | Pre-supplementation | Post-supplementation | Enrollment | Pre-supplementation | Post-supplementation | Enrolment vs Pre-supplementation | Pre-supplementation vs Post-supplementation | Enrolment vs Pre-supplementation | Pre-supplementation vs Post-supplementation |
| KEGG Functions |  |  |  |  |  |  |  |  |  |  |
| Amino Acid Metabolism | 0.098 (0.002) | 0.098 (0.0021) | 0.098 (0.0013) | 0.097 (0.0024) | 0.097 (0.0026) | 0.098 (0.0031) | 0.0379 | 0.0803 | 0.0835 | 0.0915 |
| Biosynthesis of Other Secondary Metab | 0.009 (0.0003) | 0.009 (0.0004) | 0.009 (0.0003) | 0.01 (0.0007) | 0.01 (0.0007) | 0.01 (0.0005) | 0.1333 | 0.1794 | 0.3736 | 0.2317 |
| Cancers | 0.001 (0.0001) | 0.001 (0.0001) | 0.001 (0.0001) | 0.001 (0.0001) | 0.001 (0.0001) | 0.001 (0.0001) | 0.0435 | 0.0717 | 0.1256 | 0.2135 |
| Carbohydrate Metabolism | 0.109 (0.0043) | 0.11 (0.004) | 0.109 (0.0034) | 0.11 (0.0043) | 0.11 (0.0044) | 0.108 (0.0075) | 0.2028 | 0.2562 | 0.0830 | 0.2429 |
| Cardiovascular Diseases | 0 (0) | 0 (0) | 0 (0) | 0 (0) | 0 (0) | 0 (0) | 0.1964 | 0.3691 | 0.3830 | 0.0027 |
| Cell Growth and Death | 0.005 (0.0002) | 0.005 (0.0002) | 0.005 (0.0002) | 0.005 (0.0003) | 0.005 (0.0003) | 0.005 (0.0005) | 0.1611 | 0.0653 | 0.1491 | 0.4353 |
| Cell Motility | 0.019 (0.003) | 0.019 (0.0036) | 0.019 (0.0027) | 0.018 (0.0091) | 0.02 (0.01) | 0.016 (0.0057) | 0.0145 | 0.0127 | 0.1990 | 0.5426 |
| Cellular Processes and Signaling | 0.041 (0.0008) | 0.041 (0.001) | 0.04 (0.0009) | 0.042 (0.0012) | 0.042 (0.0011) | 0.042 (0.0018) | 0.0197 | 0.0942 | 0.0184 | 0.1155 |
| Circulatory System | 0 (0) | 0 (0) | 0 (0) | 0 (0) | 0 (0) | 0 (0) | 0.1634 | 0.3326 | 0.3871 | 0.0676 |
| Digestive System | 0 (0.0002) | 0 (0.0002) | 0 (0.0002) | 0 (0.0003) | 0.001 (0.0002) | 0.001 (0.0005) | 0.1368 | 0.0681 | 0.0717 | 0.5709 |
| Endocrine System | 0.003 (0.0003) | 0.003 (0.0003) | 0.003 (0.0003) | 0.003 (0.0005) | 0.003 (0.0004) | 0.003 (0.0003) | 0.1777 | 0.0156 | 0.0244 | 0.7313 |
| Energy Metabolism | 0.058 (0.0019) | 0.058 (0.0019) | 0.058 (0.0018) | 0.06 (0.002) | 0.059 (0.0016) | 0.06 (0.0025) | 0.0926 | 0.0766 | 0.3325 | 0.5827 |
| Environmental Adaptation | 0.002 (0.0001) | 0.002 (0.0001) | 0.002 (0.0001) | 0.002 (0.0002) | 0.002 (0.0002) | 0.002 (0.0002) | 0.0488 | 0.1554 | 0.2496 | 0.8562 |
| Enzyme Families | 0.022 (0.0006) | 0.022 (0.0004) | 0.022 (0.0005) | 0.022 (0.0007) | 0.022 (0.0007) | 0.023 (0.0016) | 0.2361 | 0.1106 | 0.1057 | 0.3593 |
| Excretory System | 0 (0.0001) | 0 (0.0001) | 0 (0.0001) | 0 (0.0001) | 0 (0.0001) | 0 (0.0001) | 0.1169 | 0.0384 | 0.0349 | 0.1777 |
| Folding, Sorting and Degradation | 0.024 (0.0008) | 0.024 (0.0009) | 0.024 (0.0008) | 0.025 (0.0012) | 0.025 (0.0011) | 0.025 (0.002) | 0.0702 | 0.0680 | 0.0853 | 0.5338 |
| Genetic Information Processing | 0.027 (0.0006) | 0.026 (0.0005) | 0.027 (0.0008) | 0.026 (0.001) | 0.025 (0.0009) | 0.026 (0.0011) | 0.2298 | 0.2076 | 0.0766 | 0.2411 |
| Glycan Biosynthesis and Metabolism | 0.02 (0.0028) | 0.02 (0.0031) | 0.02 (0.0028) | 0.022 (0.005) | 0.022 (0.0031) | 0.026 (0.005) | 0.0181 | 0.0321 | 0.0454 | 0.9108 |
| Immune System | 0.001 (0) | 0.001 (0) | 0.001 (0.0001) | 0.001 (0.0001) | 0.001 (0.0001) | 0.001 (0.0001) | 0.0652 | 0.3149 | 0.1823 | 0.0025 |
| Immune System Diseases | 0 (0.0001) | 0 (0) | 0 (0.0001) | 0 (0.0001) | 0 (0) | 0.001 (0.0001) | 0.0743 | 0.0465 | 0.3842 | 0.7881 |
| Infectious Diseases | 0.003 (0.0001) | 0.003 (0.0001) | 0.003 (0.0001) | 0.003 (0.0001) | 0.003 (0.0001) | 0.004 (0.0003) | 0.0883 | 0.0247 | 0.0149 | 0.8136 |
| Lipid Metabolism | 0.029 (0.0013) | 0.029 (0.0012) | 0.029 (0.0012) | 0.029 (0.0011) | 0.028 (0.0011) | 0.028 (0.0018) | 0.1894 | 0.0815 | 0.1611 | 0.3154 |
| Membrane Transport | 0.126 (0.0103) | 0.126 (0.0094) | 0.127 (0.0097) | 0.122 (0.0138) | 0.122 (0.0106) | 0.112 (0.0171) | 0.0040 | 0.0993 | 0.0414 | 0.6641 |
| Metabolic Diseases | 0.001 (0.0001) | 0.001 (0.0001) | 0.001 (0.0001) | 0.001 (0.0001) | 0.001 (0.0001) | 0.001 (0.0001) | 0.0476 | 0.2004 | 0.1516 | 0.5964 |
| Metabolism | 0.023 (0.0003) | 0.023 (0.0004) | 0.024 (0.0004) | 0.024 (0.0005) | 0.024 (0.0006) | 0.024 (0.001) | 0.1877 | 0.1874 | 0.1942 | 0.2318 |
| Metabolism of Cofactors and Vitamins | 0.043 (0.0018) | 0.042 (0.0018) | 0.043 (0.0015) | 0.044 (0.0018) | 0.043 (0.0014) | 0.045 (0.0024) | 0.1382 | 0.0934 | 0.1551 | 0.8163 |
| Metabolism of Other Amino Acids | 0.014 (0.0005) | 0.014 (0.0006) | 0.014 (0.0004) | 0.014 (0.0008) | 0.015 (0.0007) | 0.015 (0.0007) | 0.1045 | 0.0167 | 0.3831 | 0.7379 |
| Metabolism of Terpenoids and Polyketi | 0.016 (0.0006) | 0.016 (0.0005) | 0.016 (0.0005) | 0.016 (0.0012) | 0.016 (0.0012) | 0.017 (0.0015) | 0.0044 | 0.0067 | 0.1608 | 0.6339 |
| Nervous System | 0.001 (0) | 0.001 (0) | 0.001 (0) | 0.001 (0.0001) | 0.001 (0.0001) | 0.001 (0.0001) | 0.0034 | 0.0734 | 0.1025 | 0.7113 |
| Neurodegenerative Diseases | 0.001 (0.0001) | 0.001 (0.0001) | 0.001 (0.0001) | 0.001 (0.0001) | 0.001 (0.0001) | 0.001 (0.0001) | 0.2698 | 0.1135 | 0.3445 | 0.3910 |
| Nucleotide Metabolism | 0.041 (0.0016) | 0.041 (0.0014) | 0.041 (0.0013) | 0.041 (0.0027) | 0.041 (0.0026) | 0.043 (0.0046) | 0.1118 | 0.0006 | 0.0847 | 0.5354 |
| Poorly Characterized | 0.048 (0.0003) | 0.048 (0.0006) | 0.048 (0.0004) | 0.048 (0.001) | 0.048 (0.0009) | 0.049 (0.0009) | 0.1267 | 0.2076 | 0.1446 | 0.7982 |
| Replication and Repair | 0.091 (0.003) | 0.091 (0.0024) | 0.091 (0.0026) | 0.09 (0.005) | 0.09 (0.0049) | 0.093 (0.0084) | 0.0558 | 0.1214 | 0.0152 | 0.3754 |
| Signal Transduction | 0.014 (0.0008) | 0.014 (0.0009) | 0.014 (0.0007) | 0.014 (0.0025) | 0.015 (0.0028) | 0.013 (0.0026) | 0.0939 | 0.1203 | 0.2118 | 0.5708 |
| Signaling Molecules and Interaction | 0.002 (0.0001) | 0.002 (0.0001) | 0.002 (0.0001) | 0.002 (0.0003) | 0.002 (0.0003) | 0.002 (0.0003) | 0.2380 | 0.0203 | 0.0269 | 0.9032 |
| Transcription | 0.03 (0.001) | 0.03 (0.0012) | 0.03 (0.0009) | 0.03 (0.0017) | 0.029 (0.0009) | 0.028 (0.0016) | 0.0056 | 0.0337 | 0.2626 | 0.9466 |
| Translation | 0.059 (0.002) | 0.059 (0.0017) | 0.059 (0.0017) | 0.057 (0.003) | 0.057 (0.0031) | 0.059 (0.0058) | 0.2156 | 0.0461 | 0.0095 | 0.3133 |
| Transport and Catabolism | 0.002 (0.0004) | 0.002 (0.0005) | 0.002 (0.0005) | 0.003 (0.0008) | 0.003 (0.0005) | 0.003 (0.0006) | 0.1532 | 0.0483 | 0.0104 | 0.9835 |
| Xenobiotics Biodegradation and Metab | 0.016 (0.0009) | 0.016 (0.0009) | 0.016 (0.0009) | 0.015 (0.0018) | 0.015 (0.002) | 0.015 (0.001) | 0.3328 | 0.1151 | 0.1751 | 0.1909 |
| Predicted Metagenome Mean Effect Size |  |  |  |  |  |  | 0.1182 | 0.1078 | 0.1550 | 0.4930 |

Fig 1. **Proportional abundances of dominant taxa and Shannon diversity at enrollment (EN), pre-supplementation (Pre) and post-supplementation (Post).** Panel A, Firmicutes; B, Bacteroidetes; C, Proteobacteria; D, Verrucomicrobia. Filled bars represent the OND cohort, open bars represent the OD group. Error bars depict the standard deviation for the mean of 10 subjects. p-values for significant differences are shown.

Post-supplementation, we observed a decline in Firmicutes (Fig 1A) with simultaneous increase in Bacteroidetes (Fig 1B) abundances in the OD cohort. However, only the Firmicutes were statistically different between the cohorts even though the Bacteroidetes increased by 12% over the pre-supplementation abundance in the OD group (S3 Table). The remaining taxon abundances were not significantly impacted by the supplementation but we did observe that the Proteobacteria (Fig 1C) consistently decreased, while the Verrucomicrobia (Fig 1D) consistently increased in abundance after supplementation. The abundance of individual KEGG functions were not significantly altered within or between cohorts post-supplementation (S3 Table). Alpha diversity (Shannon Index) remained significantly higher in the OND cohort at the last time point (Fig 1E).

### Changes in effect size pre- and post-supplementation

We next examined the response to the β-glucan cereal bar supplementation by comparing average effect size across the biological parameters, phyla, and predicted metagenomic content within each cohort (Fig 2, S2–S4 Tables). Firstly, effect size was not significantly different after supplementation in the OND cohort when compared to pre-supplementation abundances (p > 0.3 in all cases, Fig 2A-2C). Mean effect size was largest for the phyla (mean 0.26, range 0.04-0.62, Fig 2B), followed by the biological (mean 0.20, range 0.04-0.42, Fig 2A) and predicted metagenome content (mean 0.11, range 0.00-0.37, Fig 2C). In contrast, β-glucan supplementation had a significant effect in the OD cohort for all data categories (Fig 2A-C). Effects for the phyla and predicted metagenome content were significantly greater after supplementation while (p < 0.03, Fig 2B) while physiology effect size was significantly greater pre-supplementation for the OD (p = 0.04, Fig 2A, S3 Table).

**Fig 2.**
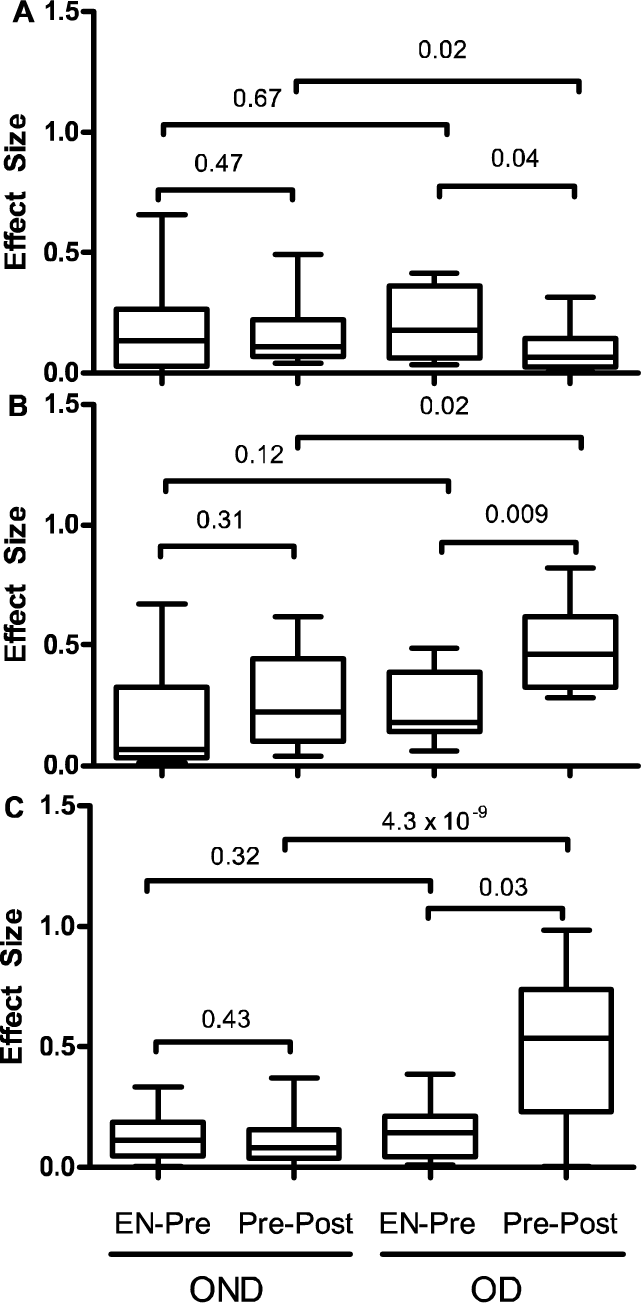
Effect size of cereal bar supplementation on measured parameters. Box and whisker plots showing the mean, the minimum and maximum values for (A) biological, (B) phyla, and (C) predicted metagenome. Kruskal-Wallis comparisons with p-values are shown. EN-Pre = effect size for enrollment and pre-supplementation time points; Pre-Post = effect size for pre-supplementation and post-supplementation time points. Data can be found in S2–S4 Tables.

Fig 2. **Effect size of cereal bar supplementation on biological (A), phyla (B), and predicted metagenome (C).** Box and whisker plots showing the mean, the minimum and maximum values for each data set. Kruskal-Wallis comparisons with p-values are shown. EN-Pre = effect size for enrollment and pre-supplementation time points; Pre-Post = effect size for pre-supplementation and post-supplementation time points. Data can be found in S2–S4 Tables.

Finally, we compared the average effect size between cohorts using the Kruskal-Wallis test to determine which cohort was more responsive to the cereal bar supplementation. Comparing the two treated groups illustrated significant differences in average effect size on the biological parameters, phyla, and predicted metagenomic content (p < 0.02, Fig 2A-C). Average effect size on biological parameters was higher in the OND compared to OD (0.19 and 0.09, respectively, Fig 2A, S2 Table), whereas average effects on the phyla and predicted metagenomic content were both higher in the OD cohorts (Fig 2B and 2C). Differences in effect sizes were most apparent for the predicted metagenomic content as there was a 4 fold greater effect in the OD versus the OND group (0.49 vs 0.11, respectively, p = 4.3 x 10^−9^, Fig 2C).

## Discussion

We tested a cereal bar supplementation containing β-glucan for 30 days in OD and OND subjects with the goal of quantifying the effect in two similar disease cohorts. We chose cereal bar with the goal of increasing fiber consumption and minimizing side effects of sudden increase of fiber intake. However, we found that consumption of 2.8 g β-glucan per day did not significantly affect blood glucose in either cohort nor did we observe improvement in any of the other individual biological parameters. The ineffectiveness of the cereal bar supplementation could result from several possibilities. For instance, the amount of fiber delivered in this study was relatively small even though it is near the FDA recommended amount for cholesterol reduction (CFR 21 101.81, [22]). Martinez et al., [17] has showed improvement in blood glucose with 60 g of additional oat fiber over four weeks in healthy individuals while DeAngelis et al. [15] showed a similar result with pasta containing β-glucan fiber in health individuals. In addition our subjects were asked not to alter their normal dietary habit and the habitual fiber consumption was not controlled, possibly confounding the effect of our fiber supplementation. We must also consider the possibility that our observations are related to the medications taken by the OD subjects as we asked the subjects not to change their drug treatment during the study. Regardless of the reasons behind our observations, we found no significant effect of 2.8 g per day β-glucan supplementation on biological parameters in our cohorts.

As we had predicted, the abundance of Bacteroidetes responded positively to the fiber supplementation in the OD cohort while there was a simultaneous decrease in the abundance of Firmicutes, suggesting small additions of β-glucan in cereal bars can alter taxonomic abundances in gut bacterial communities. These observations are in line with studies highlighting the saccharolytic nature of the Bacteroidetes in the human gut [23]. Interestingly, we did not observe any significant change in Bacteroidetes or Firmicutes abundances in the OND participants. One possible explanation is that the specific strains of Bacteroidetes and Firmicutes in the OND group are somehow resistant to change with such a small infusion of fiber. As community analysis using 16S rRNA genes does not allow for strain level resolution, an alternative approach may be necessary in order to identify specific members of the microbial community that are resistant or susceptible to cereal bar supplementations. However, it is also possible that abundances of Bacteroidetes and Firmicutes in the OND cohort are at levels where small additions of fiber can not overcome the inherent functional state of the bacterial communities requiring more intense interventions to shift the abundance of these taxa. As these two taxa represented more than 93% of the sequences in this study, the possibility exists that consuming fiber rich cereal bars may be insufficient to garner consistent and significant changes of dominant taxa across all patient cohorts.

Changes in abundance of Proteobacteria and Verrucomicrobia matched our a *priori* predictions in both cohorts. The decline of Proteobacteria is concomitant with studies indicating these bacteria are negatively associated with diets high in fiber and positively associated with high fat diets [24, 25]. The reduced abundance of the Proteobacteria has been linked to declines in inflammatory markers and general host inflammation [26]. The reduction of these organisms in human gut microbial communities is proposed to be beneficial. Conversely, increases in Verrucomicrobia are associated with healthy gut microbial communities [27] and diets high in fiber. The changes observed in both cohorts suggest that small supplements of fiber can alter abundances of these taxa. Though abundances of Proteobacteria and Verrucomicrobia did not change significantly, our data nevertheless indicates these taxa are responsive to fiber intake across similar disease cohorts possibly reflecting a general life history strategy of these organisms.

The alternative hypothesis that 2.8g β-glucan per day would have a significant impact on average effect size in both cohorts was not supported. Instead we observed significantly larger effects on the phyla and predicted metagenomic content of the OD cohort compared to the OND individuals and postulate that the lower species richness in the OD group is responsible for these observations. Greater mean effect sizes in the OD group indicate these gut communities were more susceptible to our cereal bar supplementation than were the OND group and coincide with recent report documenting richness as factor in determining microbial responses to fiber supplementations [18]. Our data build upon this observation and further suggest significantly changing microbiomes with β-glucan containing cereal bars may be limited to disease cohorts harboring microbial communities that are relatively species poor as these ecosystems have lost the functional flexibility needed to cope with a changing environment. Unfortunately our experimental design did not allow us to identify robust causal relationships between microbial communities and low intensity fiber supplementation (e.g. we used a single dosing level with small cohort size). Nonetheless, our data suggest that small amount of fiber supplementation delivered by the cereal bars have the potential to disproportionally affect low diversity microbial communities and thus may be useful in studies addressing ecological questions *in situ*, such as community stability, without altering host physiology.

Given the significant effect on the OD bacterial communities, we would have expected there to be a parallel effect on the biological parameters of these participants as gut microbes and host phenotypes are clearly linked [7, 28, 29]. However, even though effect sizes were small in both cohorts, the physiology of the OND cohort was more affected by the cereal bar supplementation. The intrinsic metabolic state of the host could explain this observation. In our subjects it may be possible that insulin resistance, pancreatic beta cell functions and even drug treatment dominate the biological set point of the diabetics and shifts in fiber consumption or bacterial communities had limited influence on the metabolic outcomes. Unlike the diabetics, these same intrinsic factors in non-diabetics may not be of equal intensity leaving a window for relatively small dietary supplementations to have some degree of influence on the host. Low dose fiber supplementation in healthy individuals reported by Martinez et al. [17] would support this observation. Altogether, the results of this study suggest bacterial community diversity in different cohorts may play a role in the response of microbes to dietary supplementation aimed to manage host metabolism.

## Materials and Methods

### Study population

A total of 46 overweight subjects with Body Mass Index (BMI) between 20 and 30 kg/m^2^ were recruited in the Service of Therapeutic education for Chronic Diseases of the University Hospitals of Geneva. After the inclusion and according to the results of the initial screening, patients were assigned to either group 1 – with type 2 diabetes mellitus (OD, n=21) or group 2 – without type 2 diabetes mellitus (OND, n=25). We defined type 2 diabetes mellitus as fasting plasma glucose > 7.0 mmol/l and/or HbA1c >7% and/or the presence of any glucose-lowering treatment. OND subjects were matched for age, gender, BMI and ethnic background of the OD subjects. Exclusion criteria were based on use of drugs altering intestinal permeability (nonsteroidal anti-inflammatory drugs, corticoids) or intestinal digestion and absorption (Orlistat, Colestipol, anticoagulants, α-glucosidase inhibitors) and antibiotics administered in the 4 weeks preceding inclusion; previous abdominal surgery, gastro-intestinal diseases interfering with intestinal absorption, cancer, bulimia, pregnancy; parenteral nutrition or other ongoing dietary intervention, and diarrhea (>2 stools/day) within 7 days before enrolment. From this initial population, 26 were excluded for compliance with the supplementation protocol, they failed to contribute samples or the sequencing effort was insufficient at any visit. The resulting subset had a mean age of 42 yrs and 51 yrs for OND and OD, respectively. Ten individuals for each cohort with overlapping age range were selected for microbial analysis. Patient data is summarized in S1 Table.

### Study design and intervention

A case controlled, single center prospective clinical trial design was used. The study consisted of 4 visits and 3 periods. The 3 periods were i. recruiting period (10 to30 days), ii. Diet-normalization period (14 days), and iii. intervention period (30 days). After their recruitment, subjects received instructions by a nutritionist for dietary normalization. After the dietary normalization period, subjects received instructions for taking cereal bars per day (one between breakfast and lunch and the other one between lunch and dinner), rich in viscous soluble fiber β-glucan. Each cereal bar contained 65 Kcal. The total carbohydrate is 10.3 g that includes 3.8 g of sugars and 2.5 g of fructose. The total protein is 1.9 g per bar. There is 1.8 g of total fat of which saturated fat is 0.7 g, monounsaturated fat is 0.7 g and polyunsaturated fat is 0.3 g. Total dietary fiber is 4.4 g of which 1.4 g is β-glucan. Sodium content is 33 mg in each bar. The cereal bar was well tolerated by all patients without clinically significant side effects. Subjects submitted blood and fecal samples at the enrollment, pre- and post- cereal bar supplementation to monitor host physiology and bacterial community composition. The study protocol (06.42NRC) was approved by the Geneva ethical committee. Participants were informed about the aims of the study and gave their written consent.

### Fecal Sample Collection, DNA extraction, PCR and sequencing

Stools were collected in sterile plastic 50 mL containers and frozen at −80°C until processing. Frozen samples were partially thawed and from which 0.25 g of fecal matter was placed in the lysis tube and extracted according to the manufacturers instructions. DNA was frozen at −20**°** C until its use in PCR reactions to generate barcoded amplicons for sequencing on the MiSeq platform [30]. Briefly, individual samples were amplified in triplicate, pooled then the PCR products were quantified using PicoGreen dsDNS reagent. Equal amounts of amplicon from each sample were then combined and sequenced on the MiSeq platform. Sequencing was performed at the Nestlé Institute of Health Sciences Functional Genomics Core facility.

### Sequence analysis

Sequence data were quality filtered and demultiplexed in Qiime 1.8 [31] using the default settings for the split_libraries_fastq.py command followed by closed reference OTU picking at 97% sequence similarity against the gg_13_5 release (http://greengenes.secondgenome.com/downloads/database/13_5) with the parallel_pick_otus_uclust_ref.py command. We additionally filtered out Cyanobacteria to avoid chloroplast sequences and filtered out low abundance OTUs according to Bokulich et al. 2013 [32] using the filter_otus_from_otu_table.py command. Samples were then rarefied to 50000 sequences per sample (single_rarefaction.py) from which the relative abundance of taxa (classified to the phylum level using summarize_taxa.py) was calculated and used for downstream analysis. An estimate of bacterial richness was performed using the Shannon index (alpha_diversity.py) on the rarefied data. Predicted metagenomic content was also calculated using PiCRUST [33] and summarized to KEGG level 2 for statistical evaluation. All bacterial community data was expressed as the proportional abundance in each sample.

### Data analysis

From the initial cohort we chose an age-matched sub set of patients to assess changes in host metabolic measurements, taxonomic abundance, and KEGG functions in response to the β-glucan cereal bar supplementation. Using Kruskal-Wallis tests implemented in Spotfire^®^ (Göteborg, Sweden), we compared group means within (e.g. pre- vs post-supplementation in each cohort) and between cohorts (e.g. OND vs OD pre- supplementation and post-supplementation) with α = 0.05 for all individual measurements (e.g. blood glucose, taxon abundance, and individual KEGG functions, etc.) to identify changes in these measurements. Bonferroni multiple comparison correction was applied to the p-values for the taxonomic abundances and KEGG functions. We next evaluated the magnitude of response of a data category (biological, phyla, and predicted metagenome; S2–S4 Tables) by testing average effect size between cohorts before and after cereal bar supplementation [34]. The absolute values of the effect size, calculated by Eq.1 where Es= effect size, m = mean, and σ = standard deviation, were compared using Kruskal-Wallis tests to determine of there were statistical differences in response.

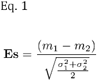

## Acknowledgements

We would like to thank Bernard Berger of the Nestlé Research Center Lausanne for his assistance, Patrick Descombes and Deborah Moine of the NIHS Functional Genomics core for their technical help with sequencing the 16S rRNA amplicons.

